# Microscope Alignment Using Real-Time Imaging FCS

**DOI:** 10.1101/2022.03.10.483720

**Authors:** Daniel Y. K. Aik, Thorsten Wohland

## Abstract

Modern EMCCD and sCMOS cameras read out fluorescence data with single-molecule sensitivity at a rate of thousands of frames per second. Exploiting these capabilities in full requires data evaluation in real-time. The direct camera-read-out tool presented here allows access to the data while the camera is recording. This provides simplified and accurate alignment procedures for Total Internal Reflection and Light Sheet Fluorescence Microscopy (TIRFM, LSFM), and simplifies and accelerates fluorescence experiments. The tool handles a range of widely used EMCCD and sCMOS cameras and uses Imaging Fluorescence Correlation Spectroscopy (Imaging FCS) for its evaluation. It is easily extendable to other camera models and other techniques and is a base for automated TIRMF and LSFM data acquisition.

**Significance:** We developed a direct camera readout (DCR) software tool that allows access to camera data during acquisition and provides real-time Imaging Fluorescence Correlation Spectroscopy (Imaging FCS) analysis. DCR displays live feedback and due to the sensitivity of correlation analysis provides a sensitive tool for microscope alignment using simple solutions of fluorescent particles. DCR is adaptable to different cameras and evaluation strategies, reduces alignment time, accelerates experiments, and can be used for automation.

## Introduction

Correlation functions are widely employed in microscopy for data evaluation, and system alignment [1,2]. They are used with static samples for microscopy alignment [3, 4, 5] and dynamic samples for the alignment in spectroscopy for instance in Fluorescence Correlation Spectroscopy (FCS) and Imaging FCS [6, 7]. Especially in the latter case, however, they are difficult to use. Because of the large amount of data acquired and the computational expensive data analysis, the steps of acquisition and analysis had to be performed in sequence. This is cumbersome as acquisition and data analysis have to be done repeatedly in iterative steps. Especially in Imaging FCS, where data acquisition and analysis can each take minutes, this is not ideal as alignment can take 15 minutes or more. During experiments, the user does not know whether a measurement is successful before all data has been acquired and treated. Therefore, there is a need for an online tool that allows real-time feedback on the quality of the alignment and data acquisition and that does not require specialised samples. We, therefore, developed a direct camera readout tool for some of the most commonly used EMCCD and sCMOS cameras that uses solutions of dyes, fluorescent beads, or supported lipid bilayers (SLBs) to provide real-time auto- and cross-correlation functions. We demonstrate that this results in quick, simple and reproducible alignment of total internal fluorescence reflection and light-sheet microscopes (TIRFM and LSM) and enables monitoring of Imaging FCS measurements during acquisition. This tool improves the alignment process for TIRFM and LSM and increases productivity, and is a simple aid for TIRFM and LSM alignment in general using solution-based samples.

## Materials and Methods

### Software

The Direct Camera Readout (DCR) data acquisition framework interfaces with a fast-acquisition camera to continuously acquire and read-out image stacks to enable users to evaluate data during acquisition at frame rates of at least 1000 frames per second. The CPU [8] and GPU [9] implementation of a multiple τ spatiotemporal correlation algorithm at the heart of DCR provides correlation functions and their figures of merit (width and amplitude) to judge the quality of data and microscope alignment. This allows real-time sample selection (Fig. S1 *A*), microscope alignment (TIRF and LSM), and long-term sample observation (Fig. S1, *B-D*). The direct camera readout routine supports multiple cameras while retaining its main features and user interface. The C++ codebase is written using software development kits (SDK) or Application Programming Interfaces (API) of the different cameras. These include SDK 2 (version: 2.103.30031.0) for the iXon DU860, iXon DU888, iXon DU897; SDK 3 (version: 3.15.20005.0) for the Sona 4.2B-11. DCAM-API (version: 20.7.6051) for the OCRA Flash4.0, PVCam (version: 2.9.0.4) for the Evolve 512 and Prime 95B cameras. The routine is included within the Imaging FCS ImageJ plugin version 1.60 (https://github.com/ImagingFCS/Imaging_FCS_1_60.git). But the code can be easily extracted and used independently or in other applications; a stand-alone java application is also made available (https://github.com/ImagingFCS/ImFCS-DirectCameraReadout.git). The simple interface design and uniform user interface enable the experimenter to focus on data collection rather than navigating complex control infrastructures associated with manufacturer-supplied software, which often are menu-driven, vary across different manufacturers, and cannot be easily integrated into user-written applications.

In this version, four routines are included. First, a *live video mode* to display live images of the sample at a screen refresh rate of 25 frames per second (fps), allowing the user to search for sample regions of interest. Second, *single capture mode* enables the user to obtain an overview of the sample and select a region of interest (ROI) for further measurements. Third, the *calibration mode* acquires ROIs and evaluates the data online, displaying spatiotemporal correlation functions for user-selected pixels at desired intervals down to every 0.1 seconds. The user can align the microscope to optimal performance by constantly monitoring the correlation functions and optimizing their width and height. Fourth, *FCS acquisition mode* acquires and simultaneously depicts the data from all pixels on an image stack, displaying the live development of the FCS measurement and providing immediate feedback to the experimenter. We have created a workflow that allows the user to screen the sample quickly, select a region of interest, and optimize alignment before finally acquiring a measurement with constant feedback on the course of the experiment.

The code is written as an ImageJ Plugin with an interactive graphical user interface (GUI) and Java Native Interface (JNI) to communicate with the C++ codebase. The program is provided in the form of the Imaging FCS software “ImFCS 1.60”, a freely available open-source plugin for Fiji/ImageJ [10] allowing routine access to any laboratory with ImageJ installed on a Windows PC with a supported camera attached to any TIRFM or LSM with no configuration required on specific cameras. Details of the plugin usage are provided in the accompanying software manual.

### Theory

FCS curves have two main parameters, their width [11,7] and amplitude, which can be used to judge image alignment. The FCS amplitude is inversely proportional to the number of particles, *N*, in the observation volume for temporal correlations. Thus a large amplitude corresponds to a low number of particles, small observation volume, and better alignment. The width of the correlation function corresponds to the average time diffusive particles need to transit through the observation volume, *τ_D_*, and smaller width corresponds to smaller observation volumes and thus better alignment. Similar arguments can be made for spatial correlations with higher amplitudes and smaller width, indicating better alignment. If we denote the *e*^-2^ radius of the observation volume by *ω*, then the FCS the amplitude for the 2D or 3D case, *G*_2*D*_ (0) or *G*_3*D*_ (0) and *τ_D_* have the following proportionalities:

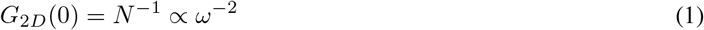

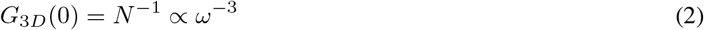

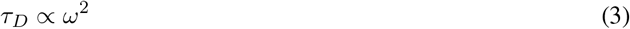

Therefore, changes in these parameters with alignment, i.e. their sensitivity, will strongly depend on *ω*:

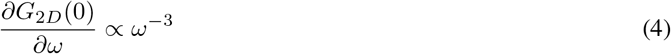

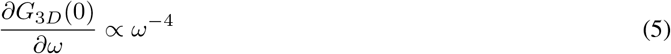

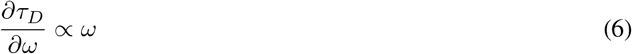

In principle, the amplitude is more sensitive to the size of the observation volume than the width. But it is also more influenced by uncorrelated background. To have a correct comparison of amplitude throughout alignment, we implemented a strategy where the minimum value within a small ROI is subtracted as a constant background across voxels before correlation calculation. This approach might not work well if strong photobleaching is observed within the small observation window (0.5 - 2 seconds). Under these circumstances, the intensity traces need to be corrected pixel-by-pixel; a unique minimum value associated with every region in space must be subtracted from the bleach-corrected trace before correlation calculation. To circumvent this problem, we instead used low laser power (80 - 160 *μ*W for the entire field of illumination) to ensure minimal photobleaching and average correlation functions over pixels in the ROI to achieve a high SNR. In the presence of background, the amplitude is given in Eq. 7 [12, 13].

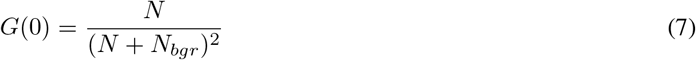

where *N* represents the number of particles in the focal volume, and *N_bgr_* the number of particles the background represents, which is given by the background signal divided by the brightness of the particles. In FCS one can distinguish two regimes (Eq. 8).

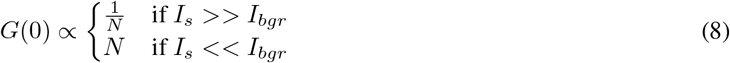

Depending on the regime in which one measures, the amplitude is not always as straightforward to interpret as compared to the width, and it is best to use both amplitude and width simultaneously to judge alignment. These two figures of merit are the basis for the alignment procedure, and the correlation function will reach a maximum in amplitude and a minimum in width when optinally aligned. The strength in comparison to other sets of evaluation criteria [4, 5]: image contrast, image variance normalized by mean, image entropy, Brenner gradient [14], and other image correlative methods [15] are that this method is highly sensitive to changes in the point spread function (PSF), can be used for alignment of the microscopes in general and is not limited to focus alignment, and does not require any structured samples but can be used with simple solutions of fluorescent particles.

### Instrumentation and Sample Preparation

Reader are referred to *Supplementary Note 1* for details in instrumentation and *Supplementary Note 2* for any sample preparation associated in article.

## Results

Many modern fluorescence microscopes come equipped with high-speed digital cameras enabling acquisition rates, at least for substantial regions of interest if not for full-chip readouts, in the range of 1000 fps for commonly available EMCCDs or 10,000 fps or more sCMOS cameras [16]. The sampling is fast enough to yield correlation functions for each pixel in the acquired image stack and thus opens the possibility to use spatial and temporal correlations for microscope alignment. Displaying these correlations in real-time provides faster and more precise instrument alignment. Furthermore, Imaging FCS based alignment requires only simple solutions of fluorescent particles and no fixed structures, making it simple and reproducible between different labs.

### Long-term Imaging FCS measurement with Direct Camera Readout

The advantage of the current program becomes evident in an SLB measurement at 1000 fps for 1,000,000 frames totalling ≈ 17 minutes measurement time. Without real-time readout and feedback, the measurement results will only be known after the full measurement time, saving of the data and offline analysis. This has to be repeated until a measurement is successful. The current program displays the development of correlation functions live, obtains correlation functions of selected ROI at user-defined time intervals and registers plots of average *D* and *N* to help the user decide whether the experiment should continue, need to be stopped, or adjustments need to be made. Furthermore, as the data is treated during the measurement, data analysis is real-time, and results are ready when the experiment stops, resulting in considerable savings of time.

Additionally, we show long term live cell observations at a full-frame EMCCD chip exposure at 2 ms frame time for close to 60 minutes, which otherwise would consume close to 60 GB of data storage. Instead, we saved the instantaneous fitted diffusion coefficient, amplitude and intensity counts of a defined ROI for illustration (Fig. S1, *B-D*). The real-time data analysis results in considerable time savings and opens up the possibility of long-term studies of cellular dynamics.

### Aligning a TIRF microscope using real-time correlations

We examined various samples ranging from solution measurements over SLBs to live cell membranes to ensure the suitability of our angle and focus alignment strategy by DCR. For this purpose, we mounted the sample on the TIRFM and then first adjusted the focus (Fig. 1 *F*) followed by the angle of illumination (Fig. 1 *A*) while observing the quality and amplitude of the temporal correlation functions (Fig. 1, *B-E, G-J*). In general, these two steps were sufficient for alignment. However, they can be performed iteratively if required (Refer to *Software manual* for flow chart). In the case of cell membranes (Fig. S2), the maximum of the obtainable correlation function coincides with image sharpness, image contrast, and the minimum width of the correlation function. This was confirmed with SLBs in which alignment by the amplitude of the temporal correlation function resulted in the highest D value, i.e. narrowest width of the correlation function, as expected [11]. This confirms that optimal correlation functions are a good measure for proper TIRFM alignment and that this technique thus does not require structured samples for microscopy alignment.

**FIGURE 1:**
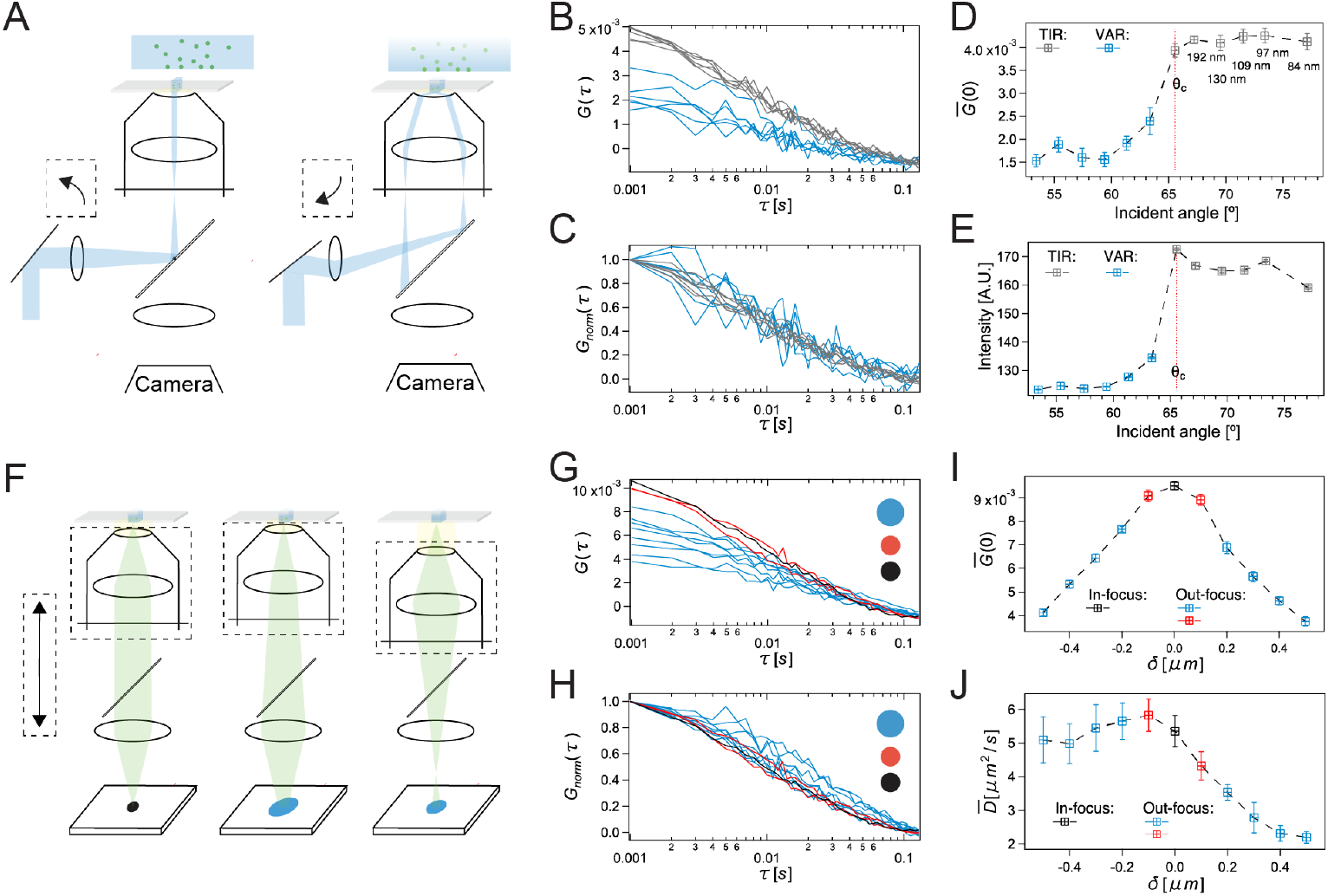
Alignment of a TIRFM system using a supported lipid bilayer with a 100 ×, NA 1.49 objective and a camera with 24 *μ*m pixel size. (A) Schematic of the TIRFM as the illumination transitions into the evanescent region at an incident angle *θ* > *θ_c_*. (B,C) Spatially averaged and normalized CFs. Each CF is calculated over 0.5 s while the incident angle is decreased stepwise without changing focus. (D, E) CF amplitude and fluorescence intensity as a function of incident angle. At the transition when *θ* > *θ_c_*, the CF amplitude, fluorescence intensity and the SNR increase. (F) Schematic of the effects of focus adjustment in the TIRFM. (G,H) Spatially averaged and normalized CFs calculated over 0.5 while the focal position is changed in steps of 100 nm. (I, J) Amplitude and width of the CF as a function of focus position. The maximum amplitude corresponds to the optimal focus (I) and is a more sensitive measure than the width of CF (J). The error bars in (D), (E), (I), (J) are calculated from 10 individual CFs, each of 0.5 s.

For TIRF measurement in solution, i.e. where the sample is in 3D instead of 2D, amplitude and diffusion coefficient peaked at a certain *z*—position and decreases monotonically in both directions (Fig. S3). The outcome of critical angle calibration (Fig. S4) for precise determination of the incidence angle and thus penetration depth is consistent with the peak evanescence intensity of immobilized beads measured on the cover glass.

The current plugin can achieve the alignment by correlation functions in under 2 minutes. But it raises the issue of the competition between sufficient SNR and bleaching during alignment. The SNR is dependent on particle brightness, and thus excitation intensity, and the square root of the number of frames [12]. To avoid photobleaching, one must minimize the excitation intensity or the number of frames acquired for the alignment. As during alignment in a solution spatial resolution for FCS is not required, we can average correlation functions of the pixels in an ROI to increase the SNR. This leads to minimal photbleaching while it maintains sufficient sensitivity for alignment.

### Aligning a Light Sheet microscope using real-time correlations

The user must align a light sheet microscope’s illumination and detection objectives with a calibration sample to ensure that the waist of the light sheet is aligned to the focal plane (alignment of the detection objective along its optical axis) and the centre of the field of view of the detection objective (alignment of the illumination objective along its optical axis) (Fig. 2 *A*) [7]. Analogous to TIRF critical angle alignment (Fig. 1), it can also be carried out in an iterative process. This is accomplished by fine-tuning the focal plane of the detection objective to the optical axis of the illumination objective and the focus point of the illumination objective to the detection volume, respectively, by maximizing the amplitude (Fig. 2 *C*), and minimizing the width of the correlation function (Fig. 2 *D*).

**FIGURE 2:**
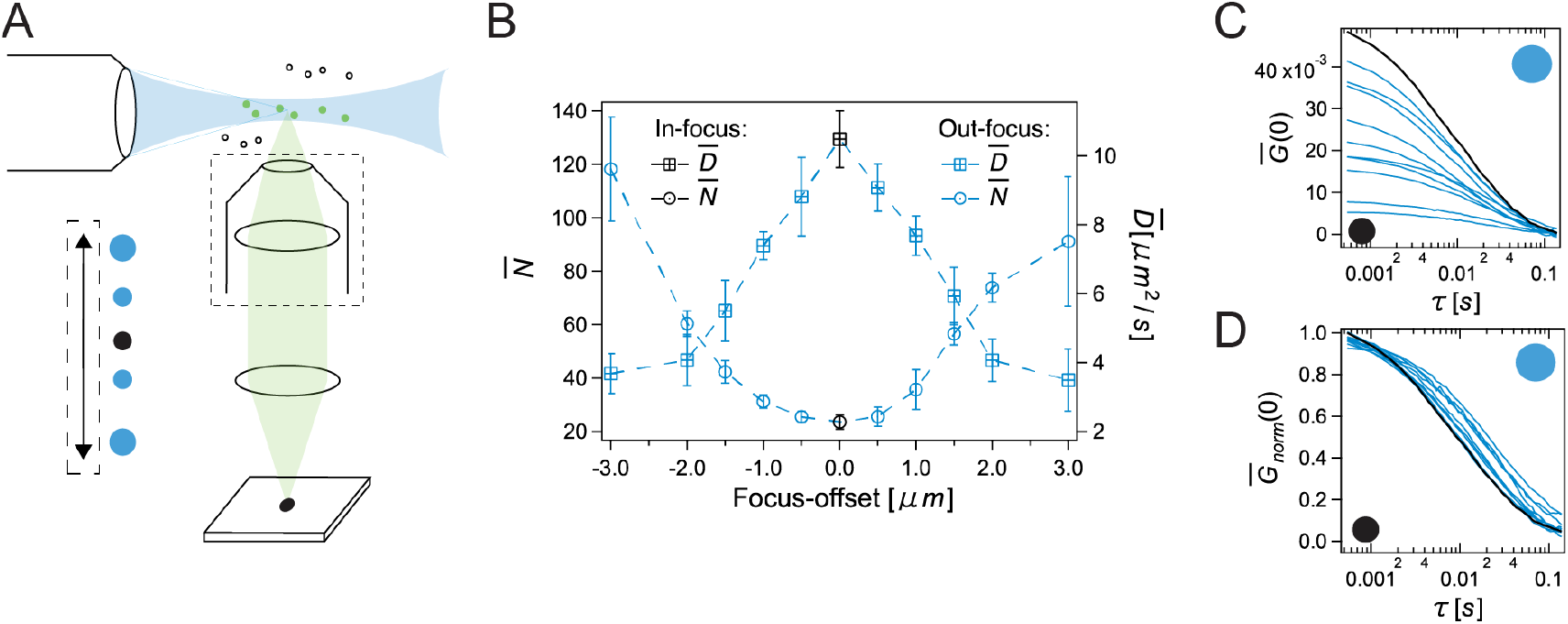
SPIM detection arm alignment using beads in solution. (B) Number of particles, *N*, and diffusion coefficient, *D*, determined from the CF at different focal positions of the detection objective. The error bars are calculated from 5 individual CFs of 0.5 s length. At optimal alignment *N* is at a minimum (largest CF amplitude) and *D* at a maximum (narrowest CF). Alignment can be achieved within 2 minutes with real-time feedback of CF, its amplitude (C) and width (D). Note that the illumination arm can be optimized similarly by altering the position of the light-sheet along the axis of propagation.

We propose a stepwise optimization starting with the detection objective, followed by the illumination objective repeated twice to ensure consistency. As the illumination and detection objectives generally differ in NA, the step sizes used for the two objectives for optimization should be chosen appropriately so that significant changes in amplitude can be observed per step size. This minimizes the time to achieve alignment. The illumination and detection arm can be treated independently; hence the order of optimization does not matter. Alternatively, one could identify a ROI associated with the thinnest light-sheet while optimizing the detection arm. Note that the latter method was used here as the light sheet is already positioned correctly along the axis of propagation. In our system of 100 nm (detection *arm)* step size leads to a clear separation in amplitude with as little as 2000 frames, with an accuracy of positioning of 500 nm for the detection objective. At 0.5 ms time resolution, 2000 frames equate to live feedback at every 1 seconds sufficient to calibrate a SPIM setup in under 2 minutes.

For calibration, by the width of the correlation function, at least 2000 frames are required, and thus correlation over the ACF amplitude sometimes is preferred. The values provided here can be used as reference but will vary among different calibration samples because sensitivity in ACF width improves with slower mobility and changes in amplitude are more noticeable with smaller effective observation volume. We recommend observing both width, amplitude, and 5 the shape of the correlation function simultaneously.

### Aligning microscopes for simultaneous dual-color experiments

In the case of dual-colour measurements, images are either spectrally separated with a dichroic mirror and images are projected onto two cameras. Or one uses an image splitter to project the two wavelength channels onto different regions of a single camera chip with (Fig. 3*A*). Here we show the alignment of a TIRF system with an image splitter. But the same alignment procedure can be used also for multiple cameras and SPIM systems. After aligning focus and angle of the TIRFM, additional steps are required to ensure the maximal overlap of observation areas between the pixels in the different wavelength images (Fig. 3 *D(top)*). Unlike single-molecule localization experiments, there is currently no solution to correct non-optimal observation areas between two channels, resulting in undesirable suppression of cross-correlation amplitude for Imaging Dual-colour fluorescence cross-correlation spectroscopy experiment [17]. We maximize the cross-correlation amplitude (Fig. 3 *C*) between two spatially equivalent detector regions by sequentially shifting either of the two images in the *x*– and *y*– direction using the image splitting device (Fig. 3*B*). To demonstrate the applicability in TIRFM, we split the images of a single fluorescent label with a 50/50 beam splitter, ensuring maximum correlation amplitude when aligned. Users can apply the same strategy with a dichroic mirror for alignment with two spectrally different fluorescent probes. So far, we have tested the proposed procedure with 110 nm to 240 nm pixel size in sample space on both EMCCD and sCMOS cameras.

**FIGURE 3:**
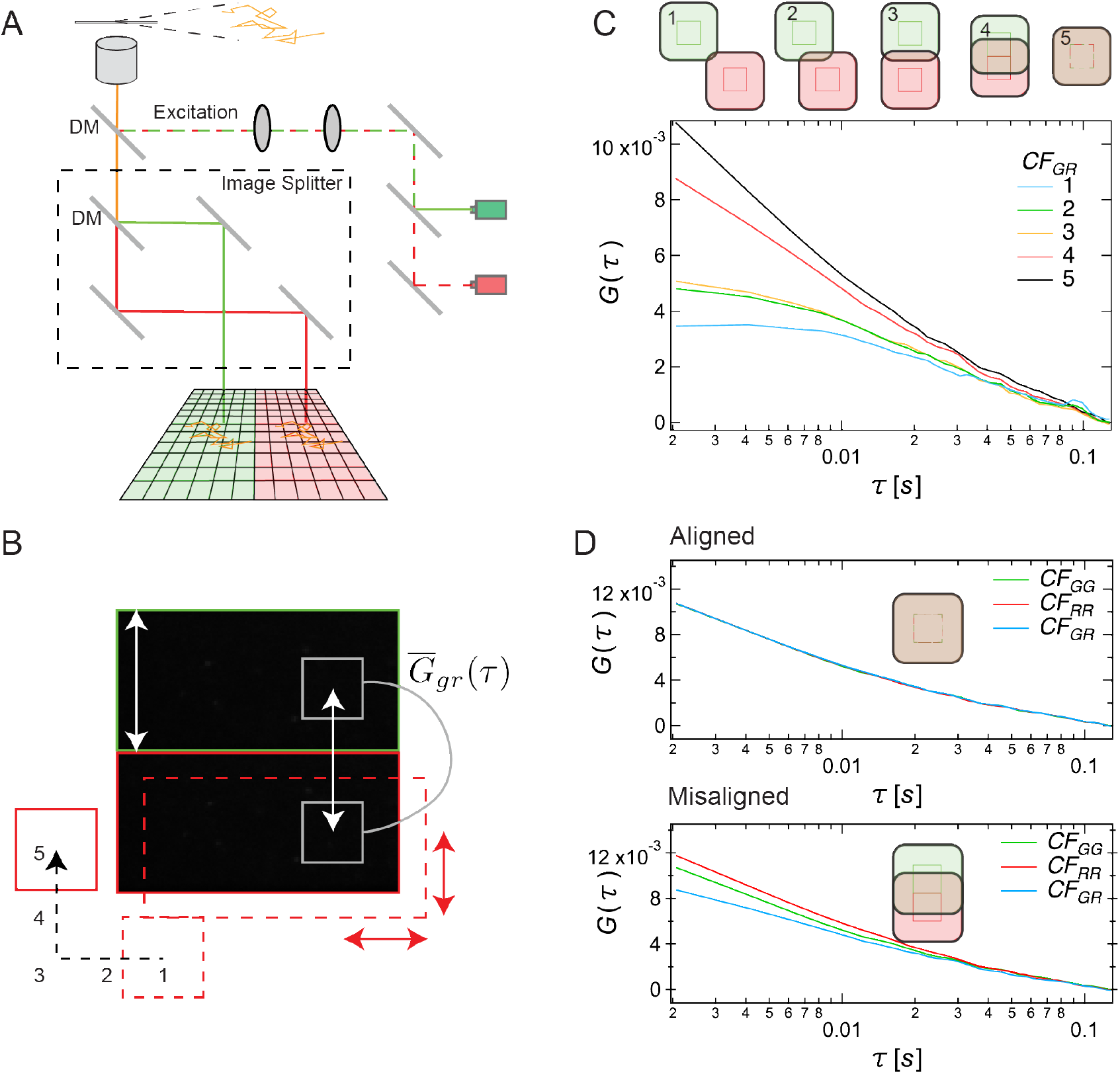
Dual-View alignment without the need of fiducial markers. (A) Emitted fluorescence light is split by wavelength (dichroic mirror), polarization (polarizing beam splitter) or intensity (beam splitter) and imaged on two halves of a camera chip, referred to as *G* and *R.* For alignment, we use a 50/50 beam splitter and a single fluorescent bead species in solution. While each pixel yields a sample characteristic autocorrelation function (*CF_GG_*, *CF_RR_*), the cross-correlations between pixels of the two halves of the chip (*CF_GR_*) depends strongly on alignment. (B) The cross-correlation function is calculated in real-time by systematically adjusting the image position of one of the channels, in our case the red channel. (C) The relation of the auto- and cross-correlation amplitudes of two pixels in the two wavelength images is a measure of the alignment. Maximal cross-correlation is reached when the pixels perfectly overlap and the amplitudes of auto- and cross-correlation functions are equal. (D) Representative CFs for an aligned system (top) and a misaligned system (bottom). Alignment can be performed directly on cellular samples (Fig. S5 *C*) with sub-pixel precision down to 48 nm for a 24 *μ*m pixel size EMCCD, using a 100 ×, NA 1.49 objective.

We compared the dynamic alignment strategy on a cellular sample using Rhodamine-PE membrane stain against a static alignment strategy on the same region using cellular structures. In addition we also used sub-resolution fiducial markers as a control (immobilized 100 nm TetraSpeck beads onto the surface of the cover glass) (Fig. S5). We calculated the spatiotemporal cross-correlation function for the dynamic sample, while for the static sample, we used the spatial cross-correlation function. We used averaging of the correlation functions over all pixels in an ROI to enhance the SNR. The alignment procedures on three different sample regions were consistent with the pixel size of the EMCCD camera used. The dynamic sample was as sensitive as the sub-resolution static sample with a reachable precision of alignment down to 48 nm (Fig. S5 *C(blue box)*). In principle one can also use cellular projections, e.g. filopodia, for alignment but is limited by the size and brightness of the structures. On cellular projections we reached a precision of only 240 nm (Fig. S5 *B(blue box))* given the setup (240 *μ*m EMCCD; 100×) and should improve in principle with smaller pixel size. This is shown by significant changes of the correlation amplitude with step size, ψ (Fig. S5 *C(middle)).* The dynamic alignment workflow performs at least as good or better than the static sample and removes the requirement of a fixed sub-resolution structure (Fig. S5 *A*). Both methods for alignment by static and dynamics samples are provided in the software.

## Conclusion

The current direct camera readout program facilitates the alignment of total internal reflection and light-sheet microscopes by using real-time calculations and display of temporal and spatial correlation functions. It uses standard bead solutions, supported lipid bilayers, or cellular samples and supports a range of commercially available cameras. With this tool, microscope alignment can be achieved within 1 - 5 minutes, depending on the type of alignment and signal-to-noise ratio. For Imaging FCS users, the program allows online monitoring of the progress of experiments, enables adjustments during measurement, and provides data evaluation in real-time, thus significantly increasing productivity. Finally, the direct camera readout can be implemented in any microscope with the equipped cameras attached. The software can be easily adapted for other camera models with supporting SDK. The alignment, acquisition and data evaluation can be automated in a motorized microscope leading to a data-acquisition pipeline with minimal human intervention.

## Software availability

The Imaging FCS 1.60 ImageJ plugin used in this paper can be downloaded in Fiji automatically followng the instructions in the manual, available here manual. Source code can be downloaded, https://github.com/ImagingFCS/Imaging_FCS_1_60.git. Direct camera readout is also available as a stand-alone sofware available here, https://github.com/ImagingFCS/ImFCS-DirectCameraReadout.git.

## Author Contributions

T.W. designed the study. D.A. programmed the algorithms and analyzed the data. T.W. and D.A. wrote the manuscript.

## Acknowledgments

We thank Ashwin V.S. Nelanuthala for help with the SPIM and Harikrushnan Balasubramanian for the construction of plasmids and live-cell sample preparation.

D.A. is supported by a National University of Singapore Industry-Relevant PhD Scholarship (NUS-IRP). TW acknowledges funding by the Singapore Ministry of Education (MOE2016-T3-1-005).

## Supplementary Information

### Supplementary Figure 1

**FIGURE S1:**
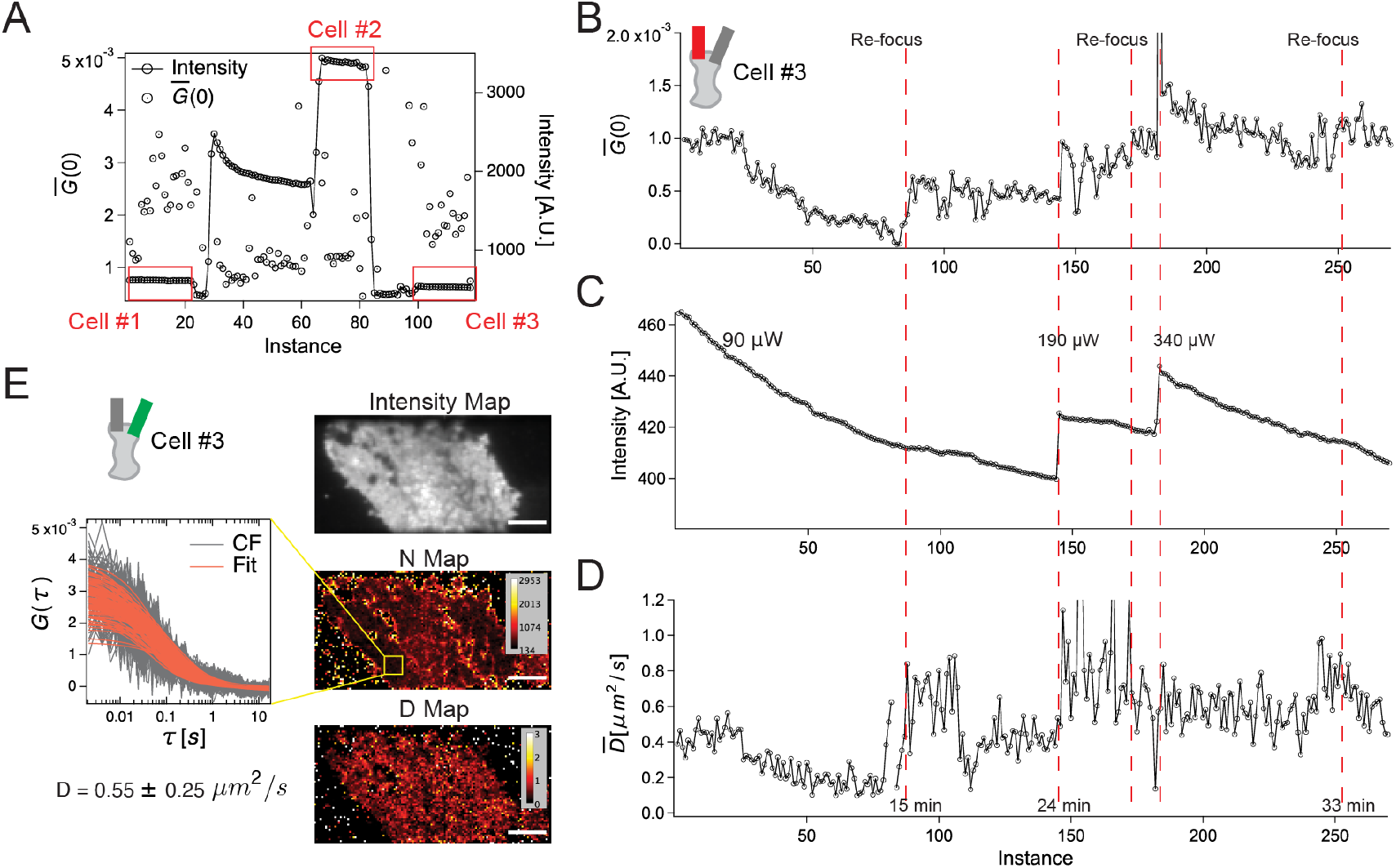
*Calibration mode* for the selection of cells expressing PMT-mEGFP-mApple and long term observation under continuous evanescent wave for over 30 min. (A) Correlation amplitude and intensity profile of potential cells while moving microscope stage. Cells with lower expression levels were selected for improved SNR. (B,C,D) Cell #3 was observed under TIR illumination for over 30 min. Focal plane and laser power were adjusted at several stages as indicated by the plot of mean intensity, G(0), and Diffusion coefficient over a user-defined ROI. The x-axis represents the updates of data in time. Data is updated every 10 seconds during observation stages and 1 to 5 seconds when refocusing. (E) A standard 100 seconds ImFCS data set acquired at the end of 30 min exposure. Intensity fluctuation from mEGFP produces diffusion coefficient maps (bottom) and number maps (middle). Scale bars represent 3 *μ*m.

### Supplementary Figure 2

**FIGURE S2:**
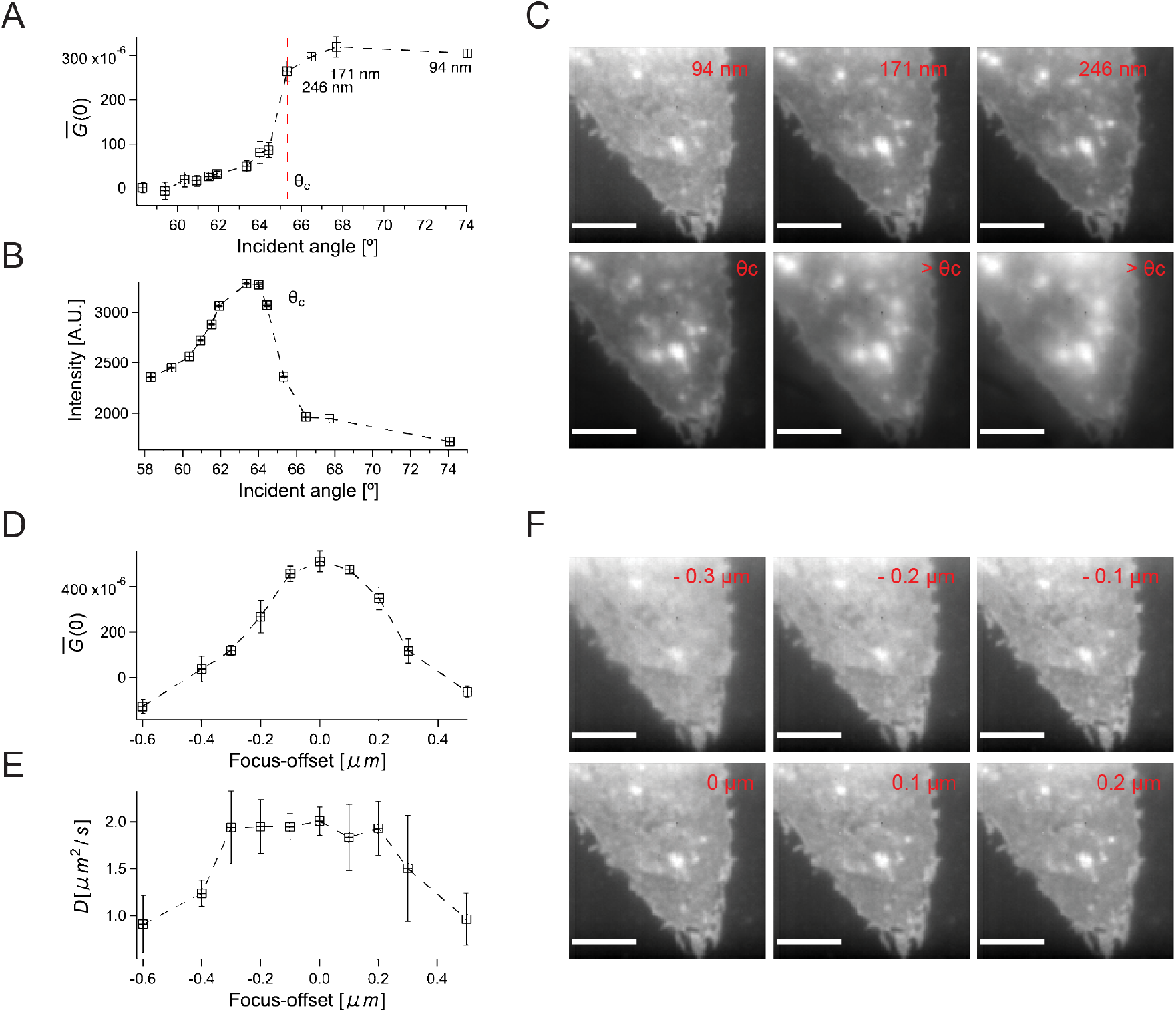
Focus and incident angle alignment of TIRFM using a standard cellular sample with a 100× magnification and an additional 2 × optical magnification. A TIRFM micrograph of a selected cell at various incident angles (C) and focus (F). A sharp increase in CF (A) amplitude indicates transition into the TIR region; therefore, the amplitude of the CF can be used as an indicator to locate the critical angle. Thus calibration can be performed directly on a sample. However, the maximum intensity (B) occurs before the critical angle, similar to TIRF in solution. The emission pattern depends on the distance of fluorophores from the surface and in a complex manner. Therefore, intensity counts cannot be used as an indicator for angle alignment due the unknown arrangement of fluorophores in the sample. The amplitude of the CF (D) is a more sensitive parameter to determine the focal plane than the width of the CF (E). The focus and angle alignment were performed with sufficiently low excitation laser power to prevent photobleaching. One can achieve focus fine-tuning in under 1 minute and angle within 4 minutes. Scale bars represent 3 *μ*m.

### Supplementary Figure 3

**FIGURE S3:**
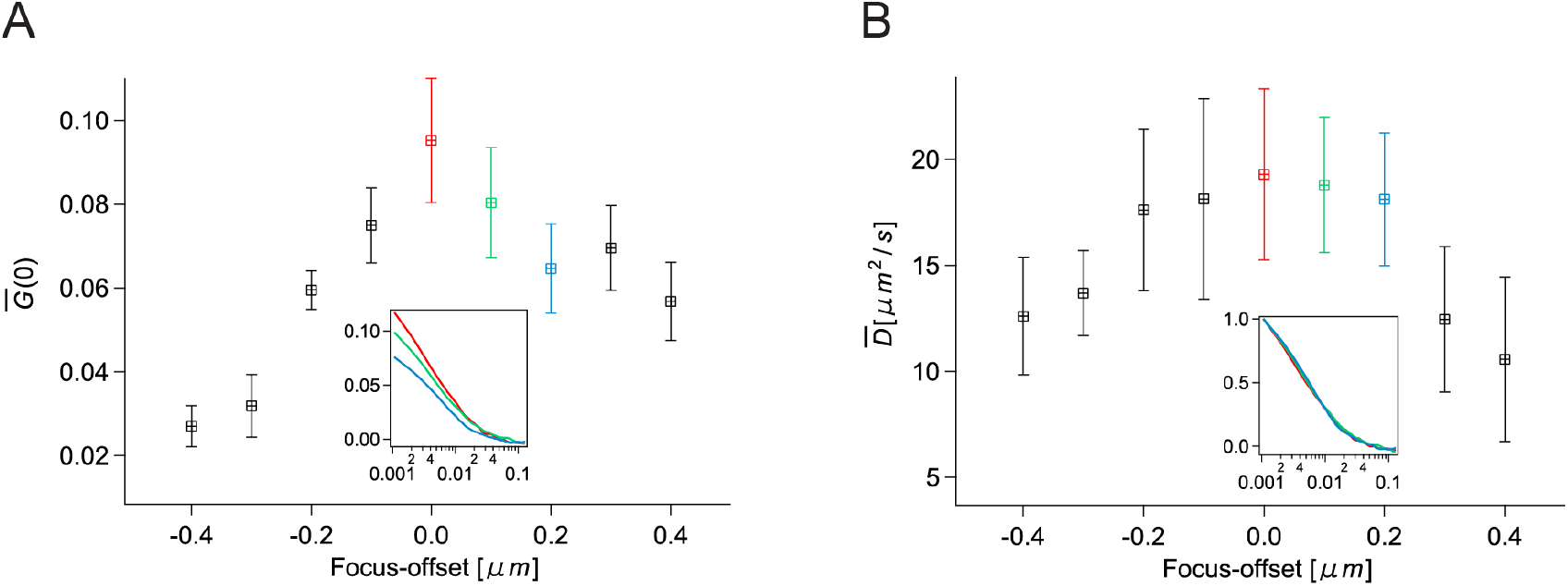
Temporal autocorrelation enables focusing TIRFM setup in solution. The amplitude G(0) and the diffusion coefficient changes as a function of z-distance for beads in solution (A, B). Data over 5 seconds were processed at a chunk of non-overlapping 500 frames to obtain the average and standard deviation. At 1 ms frame time, the user expects feedback at every 0.5 s; on average, focus fine-tuning in solution can be performed in under a minute.

### Supplementary Figure 4

**FIGURE S4:**
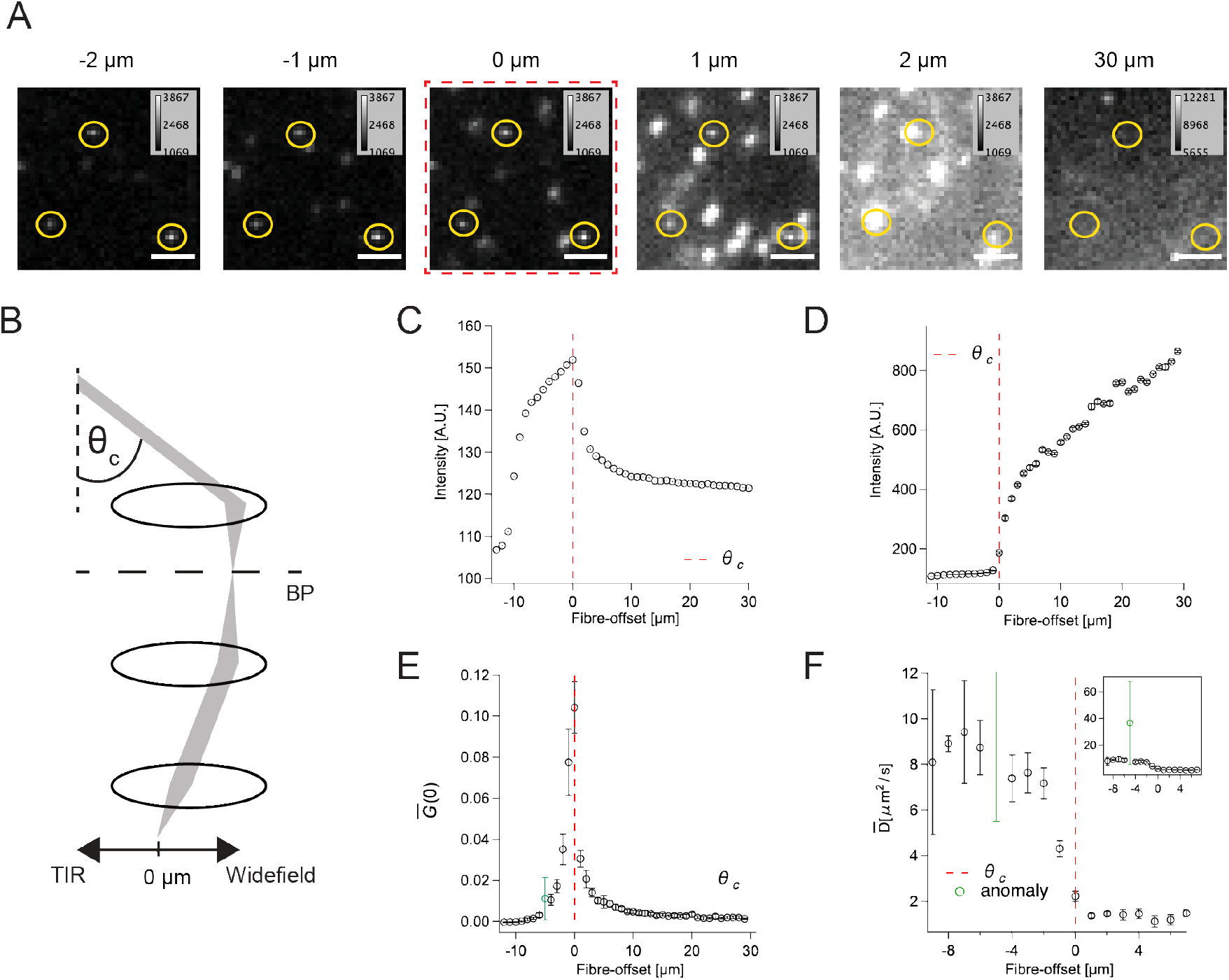
The amplitude of the correlation function G(0) is a useful metric to locate the critical angle with a dynamic sample. (A) A micrograph of beads in water at various incident angles. Immobilized beads (indicated by yellow circles) fuction as visual aid to confirm the focus. (B) Schematic of TIRF illumination. (C) Intensity, CF amplitude G(0) and diffusion coefficient of beads in solution at different angles of incidence. The fibre-position corresponding to the maximum intensity can be used as an indicator for the critical angle on immobilized beads (C). A rare outlier during the measurement is marked in green stemming from a bright, out-of-focus bead aggregate. For dynamic beads G(0) or D can be used as an indicator of the critical angle.

### Supplementary Figure 5

**FIGURE S5:**
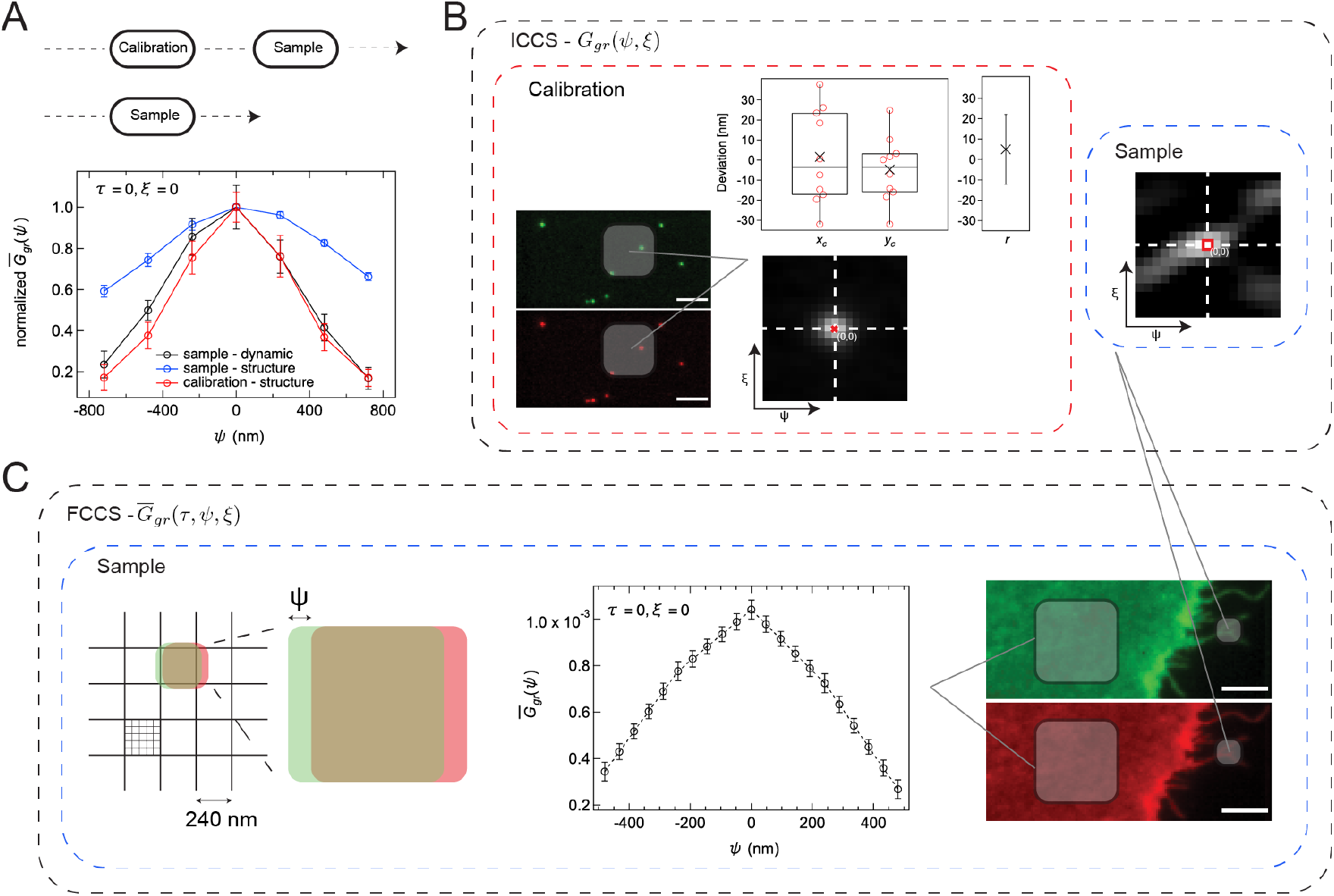
Dynamic versus static alignment strategy with sub-pixel precise alignment. (A) The results of the two alignment procedures were consistent. The dynamic sample was more sensitive, as shown by larger changes of the correlation amplitude with step size, *ψ,* for the dynamic vs the static sample. The dynamic alignment workflow performs at least as good or better than the static calibration sample, which removes the requirement of a sample with a fixed structure. (B) Image cross-correlation was performed on the sample region with visible distinct structures (blue box). The position of the maximum indicates no linear shift between two-channel, at least within pixel confidence. Composite images of sub-resolution calibration beads were analysed (red box) with the same setup. In order to validate the proposed method, CCM was computed from the selected region followed by 2D Gaussian model fitting to determine the amount of drift in each direction *x_c_, y_c_*. The Euclidean distance *r* = (5 ± 17) nm was calculated and its error propagated. In a perfectly aligned setup one expects r to be 0 nm. (C) Two spatially equivalent regions of PMT-mEGFP-mApple expressing cell were selected to demonstrate the power of temporal autocorrelation function to align an image splitter. An artificial drift was applied computationally to show the resolving power down to 48 nm. FCCS data taken at 500 frames per second with 80 *μ*W from region of 40 × 40 × 2500 were spatially averaged and plotted (middle). The error bars represent SEM. On our setup, the fiducial marker alignment strategy is precise to 40 nm corroborating the data in Figure 3. Alignment takes about 2 minutes. Scale bars represent 3 *μ*m.

### Supplementary Note 1

#### Total Internal Reflection Fluorescence Microscope (TIRFM)

Alignment of TIRFM, dual-color TIRFM, and imaging total internal reflection FCS (ITIR-FCS) was performed with two objective-type TIRFM systems. First, we used an Olympus microscope (IX83) with a motorized TIRF illumination combiner (celllTIRF-4Line IX3-MITICO, Olympus), with an oil-immersion objective (Apo N, 100×, NA 1.49, Olympus) equipped with 488 nm and 561 nm. Second, we employed a total internal reflection fluorescence microscope (IX-71, Olympus, Tokyo, Japan) with an oil-immersion objective (PlanApo, 100×, NA 1.45, Olympus) coupled with TIRF illuminator model IX2-RFAEVA-2 (Olympus), equipped with 488 nm (OBIS, Coherent, Santa Clara, CA) and 532 nm (Samba, Cobolt, AB, Sweden) lasers. The lasers are coupled into the microscope via optical fibres placed in a conjugate focal plane in both systems, ensuring collimated laser light reaches the sample. The systems have two main alignment parameters. First, the focus of the objective in respect to the sample. Second, the angle of incidence of the collimated laser light on the cover glass can be adjusted by repositioning *xy*–position of the optical fibre in the conjugate focal plane. For the dual-colour setup, we used a dual-emission image splitter (OptoSplit II, Cairn Research, Faversham, UK) placed before the camera. We used either an electron-multiplying charge-coupled device (EMCCD) or a scientific complementary metal-oxide semiconductor (sCMOS) camera of one of the following five models: 1) iXon DU860 EMCCD (24 *μ*m pixel size, 128 x 128 pixels, Andor, Oxford Instrument, UK); 2) Evolve 512 EMCCD (16 *μ*m pixel size, 512 x 512 pixels, Photometrics, Tucson, AZ); 3) Sona 4.2B-11 sCMOS (11 *μ*m pixel size, 2048 x 2048 pixels, Andor); 4) Prime 95B sCMOS (11 *μ*m pixel size, 1200 x 1200 pixels, Photometrics); 5) ORCA-Flash 4.0 sCMOS (6.5 *μ*m pixel size, 2048 x 2048 pixels, Hamamatsu, Shizuoka, Japan). Other cameras can be included in the software as long as they provide a readout speed of ≈ 1000 frames per second, and their control software is accessible.

#### Single Plane Illumination Microscope (SPIM)

The light sheet microscope used in this work is a home-built, cylindrical lens-based selective plane illumination microscope (SPIM) system with continuous illumination and detection as reported earlier [18, 19, 7, 20, 21]. We employed an identical setup according to our latest report [21]. Illumination was achieved by a 488 nm laser (06-MLD, Cobolt) coupled into a single-mode optical fibre (kineFLEX, Qioptiq, USA) which expands the beam 1.3 times, sufficient to fill the back aperture of the illumination objective. The beam from the fibre then passes through an achromatic cylindrical lens of 75 mm focal length (ACY254-075-A, Thorlabs Inc., USA) followed by an illumination objective (SLMPLN 20× /0.25, Olympus). The sample chamber of the systems is a 3 cm × 3 cm × 3 cm cube with the top open for sample mounting, glass windows on three sides made from cover glass slide thickness 1.5 to allow illumination through one of the glass windows by the illumination objective. Through a mounting hole, a water dipping detection objective (LUMPLFLN 60×/1.0, Olympus) is inserted into the sample chamber under 90° degrees to the illumination objective. The illumination objective was mounted on a manual micrometre stage (OWIS GmbH, Germany), whereas the detection objective was mounted on a piezo flexure objective scanner (P-721 PIFOC, Physik Instruments, Germany). The sample chamber is filled with water. Sample bags are made of FEP (fluorinated ethylene propylene) with a refractive index similar to water (1.338) and are lowered into the chamber. They are held using forceps placed on a motorized stage with three linear piezo *xyz* positioning systems (Q-545 Q-Motion Precision Linear Stage, Physik Instruments) and one rotation stage (DT-34 Miniature Rotation Stage, Physik Instruments). The emission obtained by the detection objective passes through a filter (FF03-525/50-25, Semrock, USA) and is projected onto an EMCCD camera (iXon DU860, Andor) by a tube lens (LU074700, f = 180 mm, Olympus). Alternatively, a flip mirror after the tube lens allows accessing a sCMOS camera (OCRA Flash4.0 V2, Hamamatsu). Time series stacks are obtained in TIFF format and subjected to the Imaging FCS analysis depending on the acquisition parameters of exposure and measurement time used. With the help of a 45° mirror, the light-sheet thickness at its thinnest point was measured to have a 1/*e*^2^ radius of approximately 1.1 *μ*m.

### Supplementary Note 2

#### Supported lipid bilayers (SLB)

All glassware (cover glass, Coplin jars, and round bottom flask) were first submerged in ten times diluted cleaning solution (Hellmanex III, Hellma Analytics, Müllheim, Germany) and sonicated for 30 minutes (Elmasonic S30H, Elma Schmidbauer GmbH, Singen, Germany) followed by 50 times rinsing with ultrapure water (Milli-Q, Merck). The glassware was then submerged in 2M H_2_SO_4_ and sonicated for 30 min followed by 50 washing steps with ultrapure water before a final round of sonication for 30 min. 0.5 mM DOPC and 50 nM Rhodamine PE (14:0 Liss Rhod PE), both supplied by Avanti Polar Lipids (Alabama, USA) were mixed in a round-bottom flask and dried in a rotary evaporator (Rotavap R-210, Buüchi, Flawil, Switzerland) for 3 - 4 h to produce a thin lipid film. The dried lipid film was rehydrated in buffer (10 mM HEPES and 150 mM NaCl, pH 7.4), resulting in a turbid solution of multilamellar vesicles before sonication until clarity to form a large unilamellar vesicle. O-rings (SYLGARD 184 Silicone Elastomer Kit, Dow, Michigan, USA) were attached to previously cleaned cover glasses (24 x 50-1, Fisher Brand Microscope, Thermo Fisher Scientific), and the above-mentioned vesicles were mixed with buffer in a 1:1 ratio and deposited onto the cover glass. The solution on the cover glass was incubated at 65 °C for 60 - 80 min followed by slow cooling to room temperature for at least 30 min. The unfused vesicles were removed by replenishing the 200 *μ*l solution inside O-rings with 200 *μ*l buffer solution at least 50 times. The resulting supported lipid bilayer (SLB) were equilibrated at room temperature and used for ITIR-FCS measurements.

#### Fluorescent beads

To remove surface contaminants, a microscope cover glass slide (24 x 50-1, Fisher Brand Microscope) was plasma cleaned (PDC-32G, Harrick Plasma, NY, USA). 100 nm TetraSpeck beads (Invitrogen, Carlsbad, CA) were sonicated to minimize aggregation and diluted by a factor of 1 to 10 for reasonable correlation amplitudes. The beads were either immobilized for static alignment on or left freely diffusing in solution for dynamic alignment on a coverslip. A 5 *μl* EtOH-beads mixture was deposited on a coverslip for immobilization. The beads on the coverslip were then submerged in ultrapure water. The solution was then dried on a heating block.

#### Plasmid

The GPI-GFP plasmid having a glycosylphosphatidylinositol-anchored protein tagged with a green fluorescent protein was a kind gift of John Dangerfield (Anovasia Pte Ltd, Singapore). The construction of the PMT-mApple plasmid consisting of a plasma membrane targeting sequence fused with mApple tag has been described in a previous publication [9]. For the construction of the PMT-mEGFP-mApple plasmid, SpeI and NheI restriction sites were added to the 5’ and 3’ terminal ends of the mApple sequence using suitable primers and amplified through a polymerase chain reaction (PCR). The PCR product was digested with Spel (Spel-HF, #R3133S, New England Biolabs, Massachusetts, USA) and HindIII (HindIII-HF, #R3104S, New England Biolabs, Massachusetts, USA) restriction enzymes. A PMT-mEGFP plasmid, similar to PMT-mApple, was digested with NheI (NheI-HF, #R3131S, New England Biolabs, Massachusetts, USA) and HindIII. The digested PMT-mEGFP and mApple PCR products were then ligated (SpeI and NheI are isocaudomers) using T4 DNA ligase (#M0202S, New England Biolabs, Massachusetts, USA).

#### Cell culture and transfection

CHO-K1 cells (CCL-61™, ATCC, Manassas, Virginia, USA) were transfected with the above-mentioned plasmids for imaging. Details on cell culture and transfection for preparation of live-cell samples are available at Protocol Exchange [22].

### Supplementary Video

It can be a challenge to align and reproduce solution measurements with TIRFM. In this video, we show how to focus and align the angle in TIRFM (**Calibration mode**) followed by 1,000 fps data acquisition with live development of ACFs for 20,000 frames (**Acquisition mode**). At the beginning of the video the TIRFM illumination is set to be above the critical angle, *θ_incidence_* > *θ_critical_*. The video starts with fine-tuning the focal position of the objective (00:26 - 00:45). The maximum amplitude (00:45 - 00:57) indicates that the objective is focused on the cover slip. Next, we change the incidence angle of the TIRFM (00:57 - 01:57) and monitor the autocorrelation amplitude. The amplitude will drop once the incidence angle decreases below the critical angle, *θ_incidence_* < *θ_critical_*. Finally, we readjust the system to *θ_incidence_* > *θ_critical_* to achieve TIR (01:57 - 02:12). A further increase of *θ_incidence_* > *θ_critical_* can reduce the penetration depth as desired (02:12 - 02:17) before data is acquired (2:45 - 3:07).

